# Lighter compared to deeper isoflurane anaesthesia can affect recognition memory in male C57BL/6 mice

**DOI:** 10.1101/645051

**Authors:** Jorge M Ferreira, Ana M Valentim

**Affiliations:** Laboratory Animal Science group, Instituto de Biologia Molecular e Celular (IBMC), Universidade do Porto, Porto, Portugal; Instituto de Investigação e Inovação em Saúde, Universidade do Porto, Porto, Portugal; Centro de Investigação e Tecnologias Agroambientais e Biológicas (CITAB), Universidade de Trás-os-Montes e Alto Douro, Vila Real, Portugal

**Keywords:** Anaesthesia, Isoflurane, Recognition memory, Mice

## Abstract

Concerns have been raised about how deeply patients are anaesthetized, and the effects that different depths of anaesthesia may have after recovery. In order to study the anaesthetic drugs per se, and to eliminate the effect of clinical variables, several animal studies have been published. Isoflurane induced transient deficits on spatial memory at low concentrations, affecting the hippocampus. However, the influence of different concentrations of isoflurane on non-spatial memory still needs clarification. Thus, our aim was to study the effects of different depths of anaesthesia (1% and 2% isoflurane) on a non-spatial memory task, the object recognition test, in C57BL/6 adult mice.

Twenty-eight 2-month-old C57BL/6 male mice were habituated to the test arena of the object recognition test for 10 min each day over 2 days before anaesthesia. Mice were then randomly allocated in different treatment groups: 1% or 2%, anaesthetized with 1% or 2% of isoflurane, respectively, for 1 h or the control group, which was not anaesthetised. Twenty-four hours after anaesthesia, the animals were placed in an arena with two identical objects and allowed to explore for 10 min-Sample Trial. One hour later, mice were allowed to explore the arena for 10 min in the presence of one of the objects presented in the previous trial (familiar object) and a novel object - Choice Trial. The time spent exploring each object was evaluated by a blinded analysis. The recognition of one object as familiar was detected based on a higher level of exploration of the novel object.

Animals that were anaesthetized previously with 2% isoflurane performed at control levels, indicating the recognition of a familiar object in the object recognition task; this contrasted with the results of the group that was anaesthetized with 1% isoflurane.

Lighter (1%) rather than deeper (2%) isoflurane anaesthesia may affect non-spatial memory in C57BL/6 male mice. Our results raise awareness of the need for careful consideration of the depth of anaesthesia used, especially the use of light isoflurane anaesthesia, which is often chosen to provide animal immobilization during non-invasive procedures.

## Introduction

Concerns have been raised regarding how deeply individuals are anaesthetized and how this depth can affect post-anaesthetic behaviour (Culley et al. 2004; Valentim et al. 2008a; Valentim et al. 2008b; Valentim et al. 2010; Zurek et al. 2012). Some studies have shown that Post-operative cognitive dysfunction was identified after light, but not deep, anaesthesia in humans (An et al. 2011; Farag et al. 2006). In mice, several studies showed that the anaesthetic that is most used in laboratory rodents, isoflurane, induces transient deficits of spatial memory (Culley et al. 2004; Zurek et al. 2012), especially when light anaesthetic depth is used (Valentim et al. 2008a; Valentim et al. 2010). However, the effect of different concentrations of isoflurane on non-spatial memory remains unclear, as different pathways are activated depending on the memory type. In this study, the object recognition test was used because it is a simple task that is applied often and it has been validated in laboratory rodents. It is a one-trial learning task of one event, thus being more sensitive to amnestic experimental interventions, such as exposure to anaesthetics (Dere et al. 2007). This task also has a good cross-species generalization (Antunes & Biala 2012), providing outcomes that can be compared with those of other species.

Thus, the objective of the study was to evaluate the effect of two different depths of isoflurane anaesthesia on the object recognition task (non-spatial task), as previous studies reported an effect of isoflurane on behavioural tasks with a spatial component in C57BL/6 male mice.

## Material and methods

All procedures were carried out under personal licenses and the project was approved (024397) by the Institutional Animal Care Committee and by the national competent authority, Direção-Geral de Alimentação e Veterinária (DGAV). Animals were housed in groups of four in Makrolon type II cages with standard corncob litter, a piece of tissue paper and a cardboard tube. The animals were housed in a room where the temperature was maintained at 21 ± 1°C and the humidity was kept at 55% on a 12 h light/dark cycle with lights on at 8 A.M. Food and water were available *ad libitum*. At the end of the experiment, animals were euthanized by cervical dislocation after the loss of consciousness induced by isoflurane.

As this was an exploratory experiment, we intended to minimize the number of animals included by using only one gender from a strain that has a well-developed exploratory behaviour, which is adequate to the object recognition task. The number of animals to be used was calculated *a priori* using the G*Power software, to achieve a meaningful statistical power of 90% and a two-side alpha error of 0.05, based on a pilot study that used untreated, healthy mice of this strain to detect significant differences (effect size of 1.7) between the exploration of familiar and novel objects. The sample-size calculation indicated that six animals were necessary. However, additional animals were used per group to avoid underpowering the experiment in case of the exclusion of animals, which was likely because the behavioural test has the requirement that each animal must explore the objects for at least 20 s; otherwise, it should be excluded. The exploratory behaviour is the central focus of this task; thus, the exclusion criterion is of utmost importance for performing the task, balancing animal behaviour and avoiding biases in the test results, to obtain comparable data among animals (Leger et al. 2013).

Twenty-eight 2-month-old C57BL/6 male mice were assigned randomly to three treatments using a computerized random-number generator (Microsoft Office Excel 2007; Microsoft, WA, USA): non-anesthetized control (n = 10), 1% isoflurane anesthesia (n = 9) and 2% isoflurane anesthesia (n = 9). A schematic representation of the experiment is presented in Figure 1A. First, mice were habituated to the empty arena of the object recognition test for 10 min each day for 2 days. In the following day, each animal was placed in an induction chamber with a gradual fill of 4% of isoflurane (Isoflo™, Esteve Farma, Carnaxide, Portugal) in 100% oxygen at 1 L/min. After righting reflex loss, the mouse was placed for 1 h in a plastic re-sealable zipper storage bag in which isoflurane concentration was maintained at 1% or 2%. The heart and respiratory rates were measured every 10 min using a pulse oximeter that was placed on the upper right hind leg of the mouse. The temperature was maintained at 37°C using a homeothermic blanket connected to a rectal thermal probe (50-7061-F; Harvard Apparatus, Ltd., Kent, United Kingdom). Isoflurane concentration was controlled in the exhausted air by an agent gas monitor (Datex Capnomac Ultima™, Helsinki, Finland), and a scavenger system was connected to the Ziploc bag. Control animals were placed in the induction chamber for 1 min to mimic the time spent conscious exhibited by mice treated with isoflurane. Subsequently, the animals were placed in their home cage, to avoid isolation stress.

**Figure 1:**
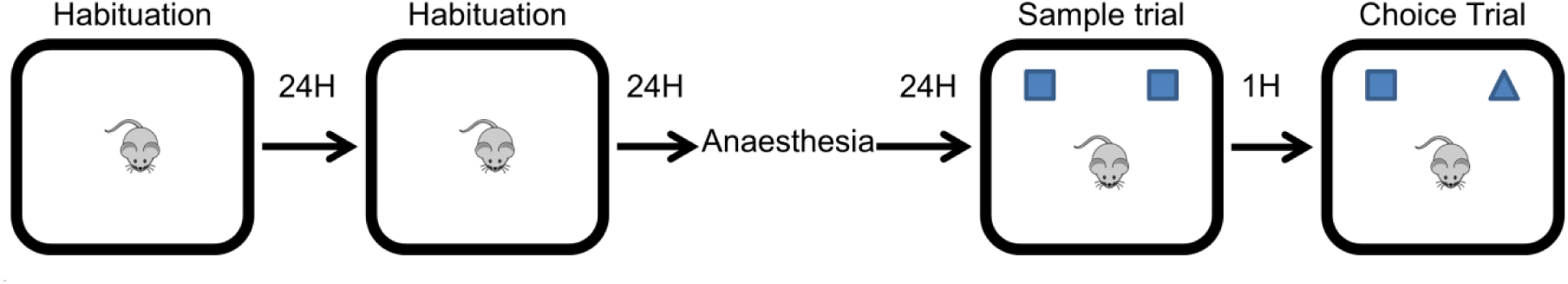
Schematic representation of the experimental design: habituation (2 days, 10 min in each day); anaesthesia took 1h; habituation, sample and choice trial are part of the object recognition task.

Twenty-four hours after anaesthesia, from 5 P.M. to 7:30 P. M., the animals were placed in the arena containing two identical objects and allowed to explore for 10 min - Sample Trial. One hour later (Inter-trial Interval), mice were allowed again to explore the arena for 10 min in the presence of one of the objects that were equal to the one presented in the previous trial (familiar object), while the other familiar object was replaced by a different one, the novel object, which was placed exactly in the same location of the familiar object that had been removed - Choice Trial (Fig. 1). Thus, there was no alteration in the spatial configuration of the arena and only one of the objects was different in the choice trial; this rendered this experiment a non-spatial test, as spatial reasoning was not required for task performance. The first 20 s of total object exploration of the choice trial were analyzed, as this is known to reduce inter-individual variability and provide a more accurate measure compared with the full duration of the trial (Leger et al. 2013). The light intensity at the centre of the object recognition arena in both trials was 400–450 lx. The behavioural task was videotaped and the time spent exploring each object was recorded by an experienced observer who was blind to the treatment using an event logging software (Observer XT, Noldus, The Netherlands) that permits the observation of the video frame-by-frame, thus allowing an accurate observation. Exploration of the object was recorded when the animal sniffed/touched the object or when its nose was within 2 cm of the object and was pointing directly to the object. This 2 cm area around each object was drawn in the computer at the beginning of the video analyses, for consistency among animals. It is considered that one object was recognized as being familiar when the time of exploration of the novel object was increased.

## Statistical analysis

Data were analyzed regarding normal distribution and homogeneity of variance between groups using the Shapiro–Wilk and Levene’s tests, respectively. Respiratory rate was analyzed by repeated-measures ANOVA with Bonferroni corrections for multiple comparisons between anaesthetic treatment and within groups (time), while heart rate was analyzed using the non-parametric test Mann–Whitney *U* test at each point of time, as this variable had a non-normal distribution. The dependent Student’s *t*-test was used to compare the duration of the exploration of the familiar object with that of the novel object during the Sample and Choice Trials. Data are presented as the mean ± S.D. All data were analyzed using IBM SPSS™ 20 for Windows (SPSS Inc., Chicago, IL, USA) after exporting it to the Microsoft Excel™ 2010 (Microsoft Corporation, Redmond, WA, USA).

## Results

During anaesthesia, mice treated with a low (1%) isoflurane concentration (n = 6 for heart rate and n=9 for respiratory rate) had higher respiratory (p< 0.0001) and heart (p≤ 0.035 from 20 to 50 min) rates compared with animals from the 2% isoflurane group (n = 7 for heart rate and n=9 for respiratory rate) (Fig. 2). There were differences in the number of individuals due to failure to acquire readings by the heart rate reader.

**Figure 2:**
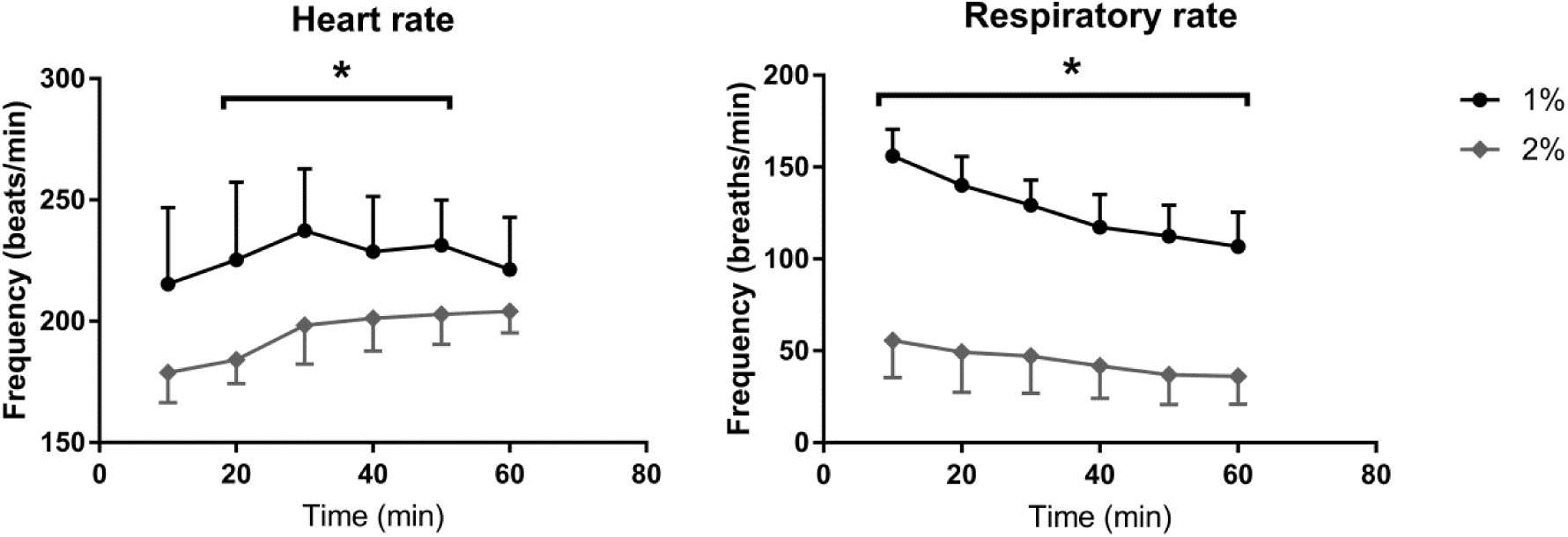
Heart rate and respiratory rate of 1% and 2% group anaesthetized during 1h. 1%: group anaesthetised with 1% isoflurane; 2%: group anaesthetised with 2% isoflurane. Data are expressed as the mean ± SD; \**p*_heart_rate_ <0.05, \**p*_respiratory_rate_ <0.0001 for comparisons between the frequency in each group (using the Mann-Whitney U test or Anova repeated measures, respectively).

In the object recognition task, three animals from the 1% group were excluded from the analysis because they explored both objects for less than 20 s during the 10 min trial (n = 6). As expected, no differences were detected in any of the groups regarding the exploration of the familiar objects during the sample trial (p_control_= 0.137, t_control_ (9) = 1.633; p_1%_ = 0. 263, t_1%_(8) = 1.205; p_2%_ = 0.799, t_2%_ (8) = 0.263). During the 20 s of total object exploration in the choice trial, control animals spent 8.3 ± 2.09 s and 11.9 ± 2.10 s exploring the familiar and the novel object, respectively. The 1% group explored the familiar object for 8.3 ± 2.89 s and the novel object for 11.7 ± 2.90 s. In the 2% group, the time spent exploring the familiar and novel objects was 7.1 ± 3.23 s and 12.89 ± 3.23 s, respectively. Thus, mice from the control and 2% group spent significantly more time exploring the novel object than the familiar one (t_control_ (9) = -1.16, p_control_ = 0.027; and t_2%_ (8) = -1.81, p_2%_ = 0.036), while no difference was detected regarding the 1% group (t (5) = -1.14, p = 0.223) during the 20 s of total object exploration (Fig. 3).

**Figure 3:**
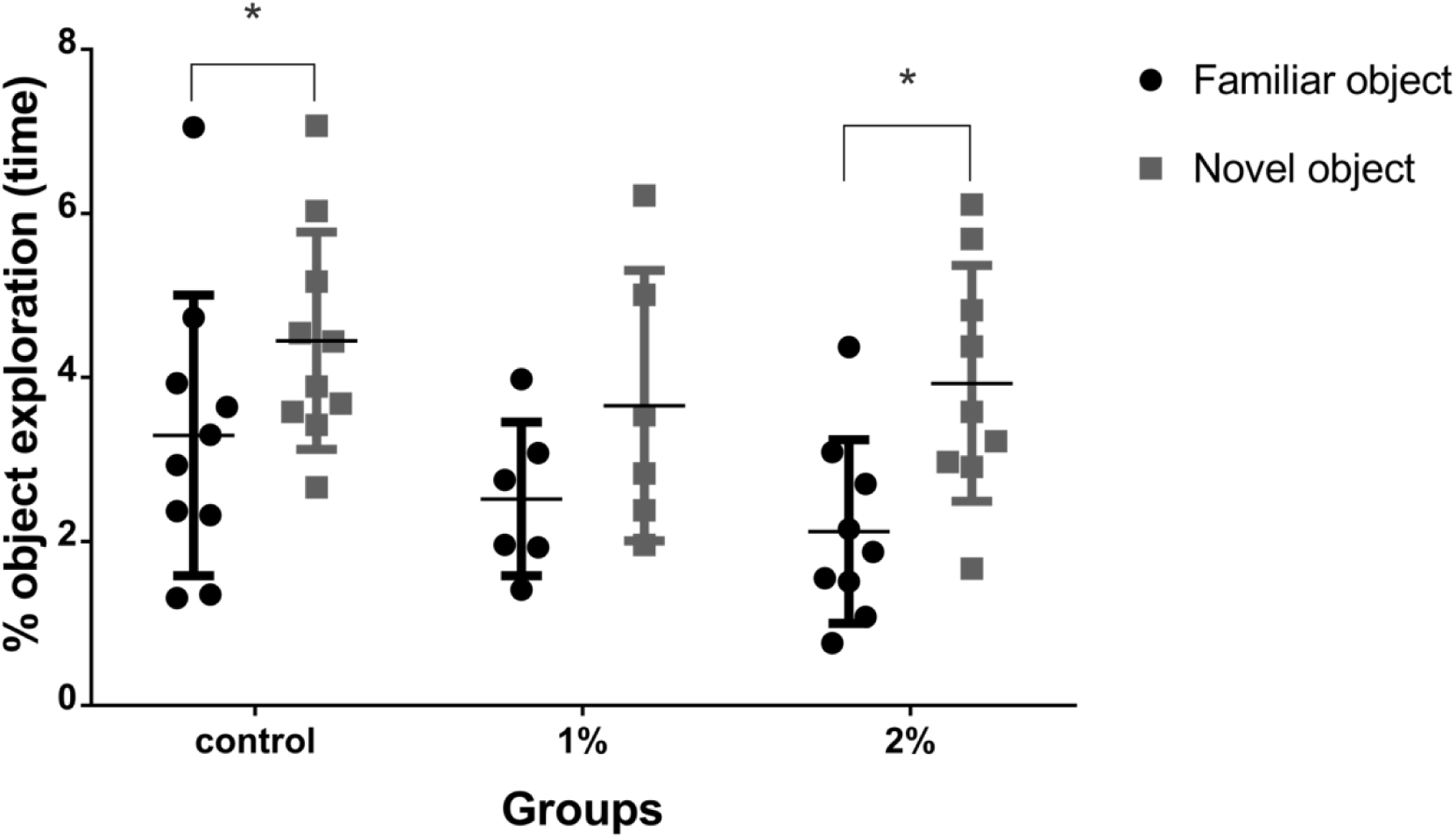
The percentage of time spent exploring the familiar and the novel object during the 20 s of total object exploration in the Choice trial of the object recognition task, 24 h post-anaesthesia. Control: group without anaesthesia; 1%: group anaesthetised with 1% isoflurane for 1 h; 2%: group anaesthetised with 2% isoflurane for 1 h. Each point represents an animal and data are expressed as the mean ± SD; \**p*_control_ = 0.027, \**p*_2%_ = 0.036 and *p*_1%_ = 0.223 for comparisons between the percentage of exploration of a familiar object and the percentage of exploration of a novel object in each group (using the dependent Student’s *t*-test).

## Discussion

Our previous works showed that a shallow anaesthesia, but not a deeper anaesthesia, induced transient deficits in cognitive spatial tasks, which can also affect the hippocampus (Valentim et al. 2008a; Valentim et al. 2010). There is evidence that non-spatial object memory in rodents is also associated with the hippocampus (Cohen et al. 2013). In fact, in this study, the 1% group seemed not to recognize the familiar object, whereas the 2% group behaved at control levels, thus revealing what appears to be an intact ability for acquiring and/or retaining information 24 h after anaesthesia. This result is partially supported by another study that reported an absence of learning impairment in the same task when the surgical concentration of isoflurane was used in adult mice (Yonezaki et al. 2015).

As we intended to understand whether different depths of anaesthesia cause different effects on a non-spatial task, we chose to compare animals treated with 1% isoflurane, a low concentration of anaesthesia (sufficient for mice to lose the righting reflex and to stay unconscious if no stimulus is applied), with animals treated with 2% isoflurane, a high concentration of anaesthesia that is able to induce surgical anaesthesia. Therefore, we were able to compare very different anaesthesia levels: 1% as the sub-MAC value and 2% as the value above MAC (MAC is 1.3% for C57BL/6 adult mice) (Sonner et al. 2000). Sub-MAC concentrations are often used to immobilize animals for non-invasive procedures such as imaging, and so the animal can be tested shortly after anaesthesia, for example, 24 h, a time at which it is fully recovered from anaesthesia but can still show side effects. In fact, our results indicate that such sub-MAC concentrations might affect recognition memory, which underscores the important consideration that anaesthesia may be a variable in research.

Other anaesthetics should be studied and used to avoid this type of behavioural side effects. For example, dexmedetomidine has been considered as an agent that does not impair memory (Zurek et al. 2012), and our group has studied it in combination with ketamine (Magalhães et al. 2017). In some situations, this combination may solve the problem addressed, or even dexmedetomidine alone may be used instead of using a low isoflurane concentration when the objective is just to calm the animals down. However, isoflurane has the advantage of providing a quick recovery and is mostly eliminated by respiration, in contrast with these injectable anaesthetics, which metabolism may interfere with other organs, such as the liver and kidney (Preckel & Bolten 2005).

As GABA-agonist anaesthetics have been shown to induce memory impairment, some authors studied the use of GABA receptor antagonists. In fact, treatment of mice with L-655,708, an α5GABA_A_ receptor-selective inverse agonist, prevented the deficits in short- and long-term contextual fear conditioning memory caused by isoflurane (Saab et al. 2010), and completely reversed the memory deficits in the novel object recognition task induced by etomidate (Zurek et al. 2012). However, the potential side effects of this reversal and its effect on the clinical efficacy of the anaesthetics should be addressed carefully.

The effects of anaesthesia described herein can vary, as they may depend on genetic polymorphisms (Behrooz 2015); thus, a larger study including additional inbred strains should be performed. We may speculate that 1% isoflurane may also decrease the exploratory motivation in mice, as the only animals excluded from our study were from the 1% group, but more studies are needed. Moreover, an intermediate anaesthetic depth should be tested to establish the dose-dependent effect and to clarify the results of Zurek et al., who showed that 1.3% isoflurane impaired object recognition performance in mice (Zurek et al. 2012).

In conclusion, we showed that exposure to 1% isoflurane anaesthesia for 1 h may impair recognition memory in male C57BL/6 mice, contrarily to 1 h exposure to 2% isoflurane. Therefore, the depth of anaesthesia should be carefully considered to attain suitable scientific quality. Our assay contributes to the existing knowledge and points to the need of further studies regarding how anaesthesia can interfere with different types of memory with different pathways of processing, to refine the data obtained from *in vivo* models.

## Acknowledgements

This work was financially supported by national funds through FCT - Fundação para a Ciência e a Tecnologia/MEC - Ministério da Educação e Ciência co-funded by FEDER funds within the partnership agreement PT2020 related with the research unit number 4293.

